# The small non-coding RNA *Vaultrc5* is dispensable to mouse development

**DOI:** 10.1101/2024.06.01.596958

**Authors:** Mahendra Prajapat, Laura Sala, Joana A. Vidigal

## Abstract

Vault RNAs (vRNAs) are evolutionarily conserved small non-coding RNAs transcribed by RNA polymerase lll. Initially described as components of the vault particle, they have since also been described as noncanonical miRNA precursors and as riboregulators of autophagy. As central molecules in these processes, vRNAs have been attributed numerous biological roles including regulation of cell proliferation and survival, response to viral infections, drug resistance, and animal development. Yet, their impact to mammalian physiology remains largely unexplored. To study vault RNAs *in vivo,* we generated a mouse line with a conditional *Vaultrc5* loss of function allele. Because *Vaultrc5* is the sole murine vRNA, this allele enables the characterization of the physiological requirements of this conserved class of small regulatory RNAs in mammals. Using this strain, we show that mice constitutively null for *Vaultrc5* are viable and histologically normal but have a slight reduction in platelet counts pointing to a potential role for vRNAs in hematopoiesis. This work paves the way for further *in vivo* characterizations of this abundant but mysterious RNA molecule. Specifically, it enables the study of the biological consequences of constitutive or lineage-specific *Vaultrc5* deletion and of the physiological requirements for an intact *Vaultrc5* during normal hematopoiesis or in response to cellular stresses such as oncogene expression, viral infection, or drug treatment.

## INTRODUCTION

Vault RNAs (vRNAs) are small non-coding RNAs initially identified in rat livers^1^ through their association with the vault complex, the largest naturally occurring cellular particle described to date^2–5^. Both vaults and vRNAs are evolutionarily conserved, and the particle itself is present at high copy numbers in many cellular contexts suggesting it has important functions in eukaryotes, though these remain to date poorly defined^6,7^.

Although initially identified via its association with the vault, only a minority of the vRNA is typically bound to that complex^8^ and many alternative functions for this RNA have been proposed. In humans, vRNAs are expressed from four related loci (*vtRNA1-1*, *vtRNA1-2*, *vtRNA1-3*, and *vtRNA2-1*) and are thought to serve as noncanonical precursors for miRNAs whose processing is DICER-dependent but DROSHA-independent^9–12^. Processing of vault RNAs into these miRNAs (also known as small-vaultRNAs; svRNAs) seems to be a regulated process^9,10^ suggesting they may act in a context-dependent manner. Studies in cell culture have suggested these svRNAs are important for proper cell differentiation^10^, required for the regulation of cell cycle progression and apoptosis^13,14^, implicated in the regulation of drug-metabolism^11^, and dysregulated in both cancer^15–17^ and neural diseases^18^. More recently, studies in human and mouse cells have shown that unprocessed vRNAs from both species can act as riboregulators of p62 to control autophagy^19,20^ a process with critical roles in cell differentiation, animal development, and human disease^21,22^. Surprisingly, despite the mounting evidence implicating both full length and processed vRNAs in the regulation of fundamental biological processes, their physiological requirements are not known.

In contrast to humans, mice have a single vRNA (*Vaultrc5* or *Mvg1*)^23^. The existence of a sole vRNA gene in murine genomes represents an invaluable opportunity to define the physiological requirements of an abundant but mysterious class of non-coding RNA molecules in mammals. With this purpose, we have generated a new mouse model carrying a conditional loss-of-functional allele for *Vaultrc5* and performed the initial characterization of *Vaultrc5*-null animals. We show that constitutive loss of *Vaultrc5 in vivo* is compatible with animal development and survival. We also find that *Vaultrc5*-null animals are histologically indistinguishable from their wild-type littermates with no detectable gene expression changes in either livers or brains, two organs where vRNAs have been proposed to play important functions. Nevertheless, we have observed a minor reduction in platelet counts in the absence of *Vaultrc5*, suggesting vRNAs may have important roles in hematopoiesis. Our work represents the first step towards defining the functions of mammalian vRNAs *in vivo*. This mouse model will enable more detailed studies using targeted and acute deletions of this conditional allele to further define the physiological roles of vRNAs during tissue development and homeostasis or in response to viral infections, oncogenic insults, or other cellular stresses.

## MATERIALS AND METHODS

### Mouse husbandry and transgenic lines

Mouse beta-actin-Cre line has been previously described ^24^ and was obtained from the Jackson Laboratory (strain 019099). *Vaultrc5* ^flx^ mice (carrying loxP sites around Vaultrc5 gene) will be made available through the Jackson Laboratory as JAX Stock No. 037602. These animals were generated by zygotic injection of a single-stranded donor DNA template ordered from IDT and *in vitro* assembled Cas9-gRNA ribonucleoprotein complexes. Reagents were generated and tested by the Genome Modification Core at Frederick National Lab for Cancer Research and used in targeting experiments by the Mouse Modeling & Cryopreservation Core. Super-ovulated C57Bl6NCr female mice were used as embryo donors. Animals were genotyped using PCR followed by Restriction Fragment Length Polymorphism (RFLP). The 3’ loxP site or the 5’ loxP were amplified using (3’LoxP, 5’-GAATCCGCGGAACTTTGG-3’, 5’-AATGCATACACAGGAGAGTTTCA-3’; 5’LoxP, 5’-AGGCAACCCATCTCTTATT-3’, 5’-GAGATGACAGACCAATCGG-3’), which amplify a 1140-bp band from (3’ LoxP) or a 1113-bp band (5’ LoxP). PCRs were cleaned up and digested with Xmnl for one hour. The resulting fragments were visualized by agarose gel electrophoresis. Correct integration in the founder male was confirmed by amplifying genomic DNA from tail clippings using primers flanking the targeting construct (5’-AGGCAACCCATCTCTTATT-3’, 5’-TGCATGTTAAAAACCCTCAGAAC-3’), cloning the resulting amplicon into the TOPO vector, followed by sanger sequencing of multiple clones. A colony for this mouse strain was established by crossing the founder to C57Bl6 females obtained from The Jackson Laboratory (Sock No: 000664). All animal procedures were conducted according to the NIH Guide for the Care and Use of Laboratory Animals, under Animal Study Proposal no. 390923 approved by relevant National Institutes of Health Animal Care and Use Committees.

### Conservation tracks

Vertebrate and Placental Mammal basewise conservation tracks were generated by PhyloP, downloaded from the UCSC genome browser, and visualized on IGV.

### RNA secondary structure predictions

Minimum free energy structure predictions were computed with RNAfold v2.4.18^25^ using the parameters ‘RNAfold -p -d2 --noLP < sequence1.fa > sequence1.out’.

### TargetScan predictions

Datasets and Perl scripts were used from 7.2 release of TargetScanMouse and TargetScanHuman^26^. First, the conserved miRNA targets and non-conserved sites were identified using a custom set of data available on TargetScan for both mouse and human^27,28^. Second, conserved branch length (sum of phylogenetic branch lengths between species that contain a site) and P_CT_ (probability of preferentially conserved targeting) for each predicted target in a custom set of data was calculated^29^. Third, the context++ scores for a set of predicted miRNA sites in a custom set of data was calculated. The context++ score for a specific site is the sum of the contribution of 14 features described in ^30^. A list of targets with predicted miRNA binding sites in their 3’ UTRs was generated based on conserved branching length, P_CT_ and context++ score. The entire analysis was done on *Homo sapiens*, *Mus musculus*, *Macaca mulatta*, *Pan troglodytes*, *Bos taurus*, *Rattus norvegicus* and *Monodelphis domestica* which were available on TargetScan.

### Histology and blood collection

Eosin-Hematoxylin staining was performed on 5 μm sections of tissues fixed in 10% neutral buffered formalin and embedded in paraffin. Blood was collected by cardiac puncture in anesthetized animals. Animals were euthanized immediately upon completion of blood collection in K2-EDTA collection tubes (Thomas scientific). Complete blood counts as well as histopathology analysis were performed at MD biosciences.

### Cell lines and cell culture

Mouse Embryonic Fibroblasts (MEFs) were derived using standard protocols. Briefly, embryos were isolated from *Vaultrc5*^+/-^ intercrosses at embryonic day (E) 13.5. After removal of internal organs, embryo carcasses were minced and digested with trypsin at 37°C before enzyme inactivation with DMEM media (GIBCO) supplemented with 10% FBS, penicillin/streptomycin (100 U/mL), and L-glutamine (2ITmM). For each embryo, the resulting cell suspension was plated on a 10 cm dish. Once confluent, these primary cells were frozen as passage 1. To generate immortalized cell lines, early passage primary MEFs from all genotypes were infected in parallel with the SV40 large T antigen (Addgene:13970) (Zhao et al., 2003). MEFs were maintained at 37°C (5% CO_2_) in DMEM media (GIBCO) supplemented with 10% FBS, penicillin/streptomycin (100 U/mL), and L-glutamine (2ITmM).

### RNA isolation and quantitative RT-PCR

Quantitative real time RT-PCR (qPCR) was performed on RNA isolated using TRIzol reagent (Invitrogen) using standard methods. RNA was treated with ezDNase (ThermoFisher Scientific) and reverse transcribed using SuperScript IV Reverse Transcriptase (Thermo Fisher Scientific) using random hexamer primers. Relative expression of *Vaultrc5* and *Gapdh* was quantified using using Sybr Green (ThermoFisher Scientific) and previously validated primer sets (Bodak et al., 2017) (*Vaultrc5*, 5’-AGCTCAGCGGTTACTTCGAC-3’, 5’-TCGAACCAAACACTCACGGG-3’; *Gapdh* 5’-AGGTCGGTGTGAACGGATTTG, 5’-TGTAGACCATGTAGTTGAGGTCA-3’).

### Small RNA sequencing analysis

Small RNA seq data for different mouse tissues were downloaded from GEO DataSets (GSE119661)^31^. Adapter sequence TGGAATTCTCGGGTGCCAAGG was trimmed using parameters; -e 0.1 -n 1 -O 3 -q 20 -m 15 -M 25. Trimmed reads were mapped to customed non-coding RNA genome (Mus_musculus.GRCm39.ncrna) using STAR aligner (2.7.11b) with --outFilterMismatchNmax 1 \ -- outFilterMismatchNoverLmax 1 \ --outFilterMismatchNoverReadLmax 1 \ --outFilterMatchNmin 16 \ -- outFilterMatchNminOverLread 0 \ --outFilterScoreMinOverLread 0 \ --outFilterMultimapNmax 5000 \ -- winAnchorMultimapNmax 5000 \ --seedSearchStartLmax 30 \ --alignTranscriptsPerReadNmax 30000 \ --alignWindowsPerReadNmax 30000 \ --alignTranscriptsPerWindowNmax 300 \ --seedPerReadNmax 3000 \ --seedPerWindowNmax 300 \ --seedNoneLociPerWindow 1000 \ --outFilterMultimapScoreRange 0 \ --alignIntronMax 1 \ --alignSJDBoverhangMin 999999999999. Bam and Bigwig files were generated using samtools (v 1.19) and exported to Integrative Genomics Viewer (2.16.2) to visualize reads to the Vaultrc5 and mmu_miR-19a loci. Expression of svRNAs and mmu_miR-19a were represented in Reads Per Million (RPM).

### Total RNA sequencing

RNA from brains and livers from *Vaultrc5*^+/+^ and *Vaultrc5*^-/-^ mice was isolated using TRIzol reagent (Invitrogen) following the manufacturers’ protocol. RNA was then treated with DNase I using Zymo RNA clean & concentrator kit. Libraries were constructed using the Illumina® Stranded Total RNA Prep, Ligation with Ribo-Zero Plus and sequenced with stranded sequencing on NextSeq2000. Reads were trimmed using cutadapt (v 4.7) for all Illumina adapters with parameters; --nextseq-trim=20 --trim-n -n 5 -e 0.1 -O 3 -q 20,20 -m 25 then trimmed reads were mapped to Mus_musculus. GRCm39 genome with Ensemble Genes 112 using --outFilterMultimapNmax 25 \ --winAnchorMultimapNmax 50 \ -- alignTranscriptsPerReadNmax 10000 \ --alignWindowsPerReadNmax 10000 \ -- alignTranscriptsPerWindowNmax 100 \ --seedPerReadNmax 1000 \ --seedPerWindowNmax 50 \ -- seedNoneLociPerWindow 10 \ --outFilterMultimapScoreRange 1 \ --outFilterMismatchNoverLmax 0.3 \ --sjdbScore 2 using STAR aligner (2.7.11b). Read alignments were counted using Subread featureCounts (v 2.0.3) with -s 0. Differentially expressed genes were analyzed by using DEseq2 (R 3.6.0).

### Statistical analysis

Statistics were done using R v4.1 (Team, 2018).

### Data availability

RNA sequencing datasets have been deposited to GEO and are available under GSE269048.

## RESULTS

### Analysis of non-canonical miRNAs derived from the murine *Vaultrc5* locus

Mouse genomes encode a sole vault RNA gene, *Vaultrc5* (also known as *Mvg1*)^32^. *Vaultrc5* is transcribed as a single exon-transcript from chromosome 18 (18qB2) from a locus located immediately downstream of *Zmat2* (**Supplementary Figure 1A**) and that is in a syntenic region conserved across mammals^23^. All vRNAs are characterized by their ability to associate with the Vault particle, are often transcribed from analogous genomic locations, and have been described in eukaryotes ranging from amoebas to humans. However, the *Vaultrc5* locus shows relatively low sequence conservation to other vertebrates or placental mammals when compared to that of protein coding genes (**Supplementary Figure 1A**). Nevertheless, a few features of the locus show some conservation. First, like vault RNAs of other species, the regulatory elements in its polymerase III (pol III) promoter are well defined (**Supplementary Figure 1B**). These include a proximal element and a TATA-like box^32^. Both of these are also found in the promoters of human *vtRNA1* genes (*vtRNA1-1*, *vtRNA1-2*, *vtRNA1-3*)^32^, which like *Vaultrc5* are located downstream of *Zmat2*. Moreover, at least the proximal element shows broad conservation amongst placental mammals and vertebrates as does the polymerase III termination sequence (**Supplementary Figure 1B**) (see also ^23^). Within the gene body itself, *Vaultrc5* contains three regulatory sequences in the form of the A and B boxes^5^, which characterize pol III type 2 promoters such as those found in tRNA genes^33^. Only two of these (A box and B2 box) are broadly conserved, as are the sequences that surround them (**Supplementary Figure 1B**). In contrast, the central region of vRNAs varies substantially between species^23^. In the case of *Vaultrc5*, it contains a third pol III internal sequence known as B1 box, with low overall conservation with other species (**Supplementary Figure 1B**) with the notable exception of rats^32^.

Despite these differences, the 3’ and 5’ ends of *Vaultrc5*, are predicted to form a double stranded region that is identical to that predicted for human *vtRNA1* genes, as exemplified here by *vtRNA1-1* (**Figure 1A**). Importantly, although the sequences predicted to fold into this double stranded structure are highly conserved between vRNAs of the two species (**Supplementary Figure 1C**), they share only 83- 89% of nucleotide identity. Yet, every position at which the sequence of the mouse *Vaultrc5* differs from that of one of more *vtRNA1* gene, the change either has little impact on base pairing (e.g. G·C pairing between bases 3 and 136 of *Vaultrc5* versus G·U wobble pairing between bases 3 and 92 in *vtRNA1-1*) or is compensated by changes in the complementary RNA sequence (**Supplementary Figure 1C**, **Figure 1A**). This suggests that the secondary structure of the stem may play important roles for the functions of vRNAs, and that those are conserved between mice and humans. Given that for human *vRNA1-1* this stem is processed by DICER into non-canonical miRNAs^11^ (**Figure 1B**), we thought it was likely that identical molecules are produced from *Vaultrc5* as well (**Supplementary Figure 2A**). To investigate this possibility, we analyzed small RNA sequencing datasets from a variety of adult mouse tissues^31^. We found that reads matching the predicted location of the mature miRNA sequences were relatively low abundant (**Figure 1C, 1D, Supplementary Figure 2 B-D**). Of all four predicted non-canonical miRNAs derived from this locus, *svRNA\** had the highest number of reads. Nonetheless, they were much less abundant than those matching canonical miRNAs. as an example, *miR-19a-3p* a miRNA with functions in hematopoietic tissues reached about 9,000 RPM in bone marrow and spleen samples, with a mean of around 2,600 RPM across all tissues (**Figure 1D**). In contrast, reads for the predicted *vaultRNA* -derived miRNA-like fragments had an abundance similar to that of *miR-19a-5p*, the passenger strand of miR-19a which is degraded following loading of the mature miRNA into an Argonaute protein (*miR-19a-5p* max=39.3 RPM, mean=6.3 RPM; *svRNAa* max=5.2 RPM, mean=0.7 RPM; *svRNAa\** max=130.8 RPM, mean=20.7; *svRNAb* max=3.8 RPM, mean=0.5 RPM; *svRNAb\** max=3.8 RPM, mean=0.5 RPM). Low counts were observed even in lung, where the levels of *vaultRNA* -derived miRNAs were reported to be high in humans (mean counts *svRNAa* =0.4 RPM, *svRNAa\** =25.5 RPM, *svRNAb* =0.2 RPM, *svRNAb\** =0.2 RPM)^11^. Given that the strength of repression imparted by a miRNA depends strongly on its abundance^34^ and that even well-expressed miRNAs like *miR-19a* have typically only modest effects on target gene expression these data suggest that under homeostatic conditions *Vaultrc5* is not a source of non-canonical miRNAs with physiologically relevant functions in the tissues we have analyzed.

**Figure 1.**
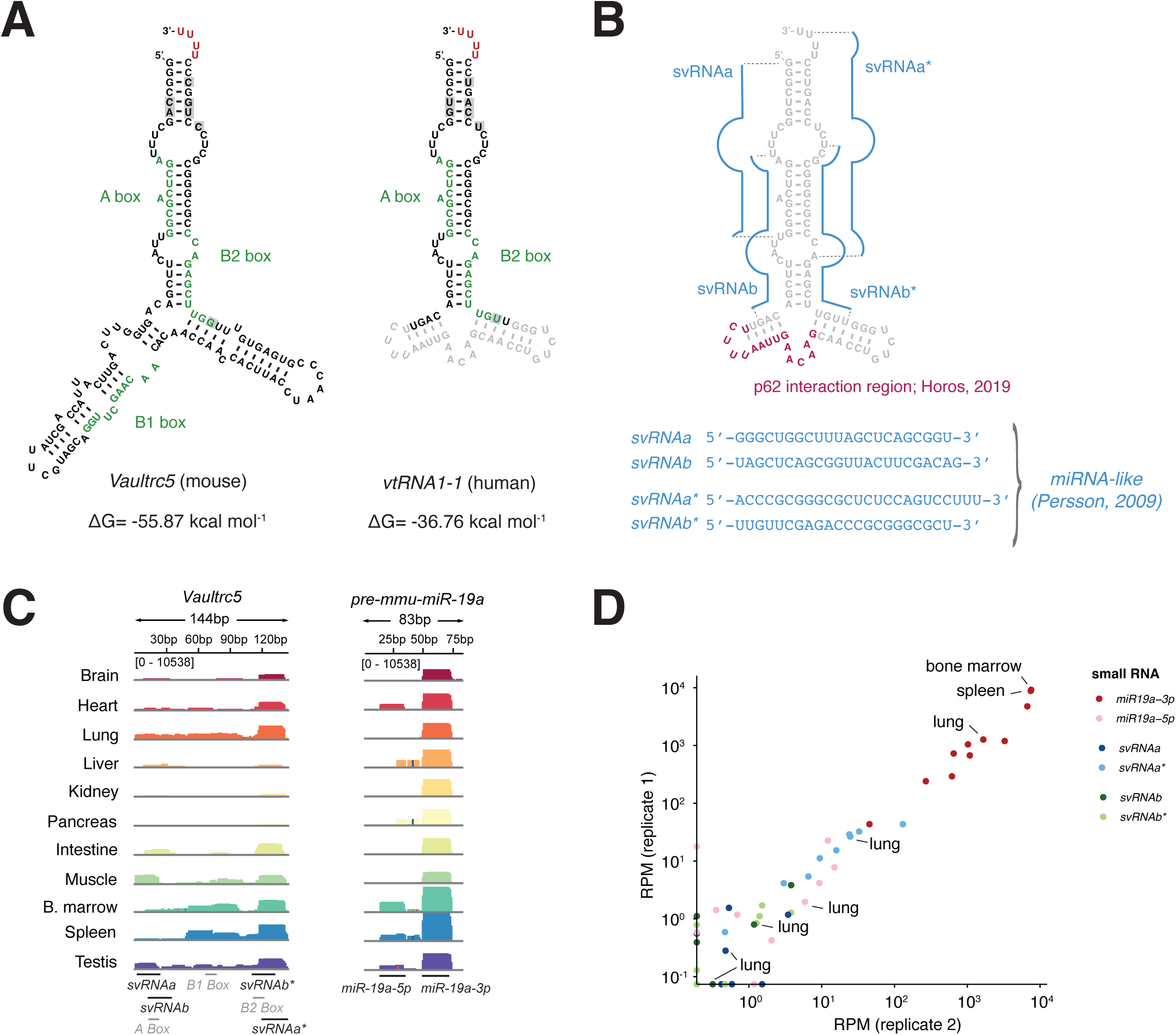
Structure and conservation of the murine *Vaultrc5* locus. **(A)** Minimum free energy secondary structure prediction for *Vaultrc5* and *vtRNA1-1* transcripts. The position of the internal regulatory elements is highlighted. The sequences that are not conserved between the mouse and human transcript are shown in grey in the *vtRNA1-1* structure. **(B)** Position of known functional elements on *vtRNA1-1* including the non-canonical miRNAs (blue, sequences shown on the bottom) and p62 interaction region (magenta). **(C)** Genome browser view of sequencing reads over *Vaultrc5* and *pre-mmu-miR-19a* loci in multiple adult mouse tissues **(D)** Expression levels of the predicted Vaultrc5-derived miRNAs compared to those *of miR-19a* (*miR-19a-3p*) and its passenger strand (*mir-19a-5p*) across two sequencing replicates.

Previous work has suggested that in humans, *svRNAb* plays a role in drug resistance through the regulation of target genes such as *CYP3A4* ^11^. Based on this, it is possible that *vRNA* -derived miRNAs perform conserved regulatory functions only under stress conditions. We think that is unlikely to be the case. First, miRNAs recognize their targets primarily through base-pair complementarity via their extended seed regions (miRNA nucleotides 2-8)^35^. Although the sequence variations between the *vtRNA-1* and *Vaultrc5* genes preserve the secondary structure of the *vaultRNA* stem, they do not preserve the seed of all predicted svRNAs, specifically *svRNAa* and *svRNAb\** (**Supplementary Figure 2A**). Second, although the predicted seed sequences of *svRNAb* and *svRNAa\** is shared between the two species, there is poor overlap between their predicted targets, based on our own implementation of the targetscan algorithm^26^ (**Supplementary Table 1**).

### Generation of a *Vautrc5* conditional loss-of-function allele

Despite the lack of evidence that *Vaultrc5* functions as a non-canonical precursor for murine miRNAs, the conserved secondary structure of its stem across species suggests this non-coding RNA plays important functions *in vivo*, potentially as a component of the Vault particle or in riboregulatory role of p62. In fact, Vault RNAs have been functionally implicated in processes such as cell survival^36^, proliferation^37^, and differentiation^10,38^, all of which are essential for successful embryonic development. The expression of full length vRNAs also seems to be regulated during cellular differentiation^38^. Together these reports suggest that full-length vault RNAs may be essential during mammalian embryogenesis. To help define what those functions might be we generated a conditional loss-of-function allele that would allow both constitutive as well as spatiotemporally controlled deletion of the locus enabling the dissection of the physiological requirements for *Vaultrc5* expression in mice. We designed a construct in which *loxP* sites were inserted directly upstream and downstream of the annotated promoter and termination elements respectively without disrupting them (**Figure 2A**, **Supplementary Figure 1B**). Both *loxP* sequences were inserted at genomic sites with no sequence conservation amongst vertebrates or placental mammals suggesting they do not unintentionally disrupt regulatory elements of neighboring genes. Moreover, no conserved sequences aside from those of *Vaultrc5* are predicted to be affected by Cre-mediated recombination of the *loxP* sites. Of note, putative enhancer elements as well as CTCF binding sites are found in the vicinity of the *Vaultrc5* but, as above, none are affected by our targeting or by the subsequent CRE-mediated genomic deletion. To facilitate the genotyping of the *Vaultrc5* locus following targeting we included an *XmnI* restriction site next to each *loxP* sequence (**Supplementary Figure 1D**).

**Figure 2.**
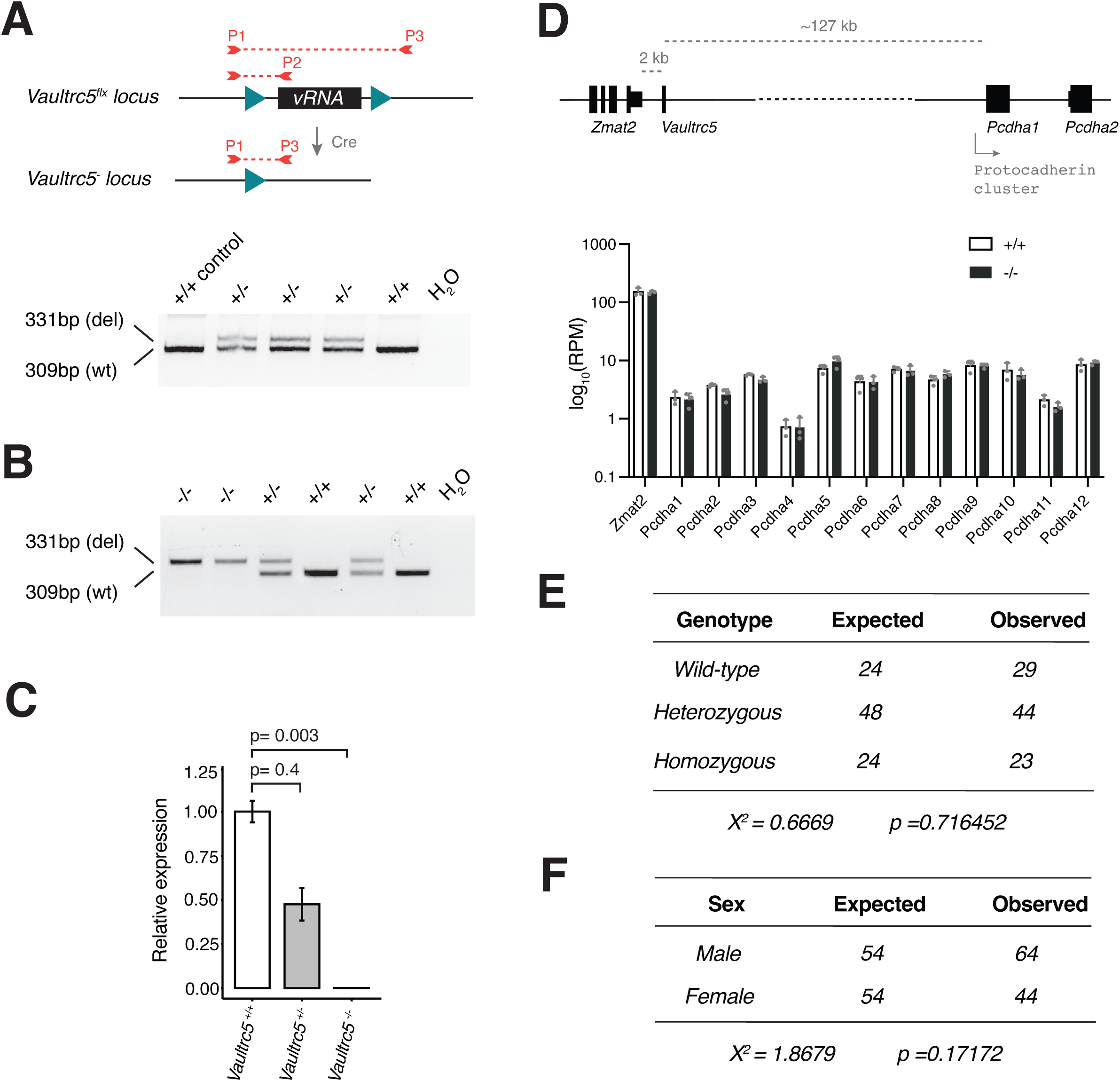
A conditional loss of function allele for *Vaultrc5*. **(A)** Schematic representation of conditional *Vaultrc5* locus (*Vaultrc5^flx^*) and its recombination after Cre expression to generate a null allele. Position of the three primers used for monitoring the recombination is highlighted. When used in combination these result in amplicons of 309 base-pairs (bp) for wild-type alleles (P1/P2) and of 331 bp for recombined alleles (P1/P3). An example gel is shown on the bottom showing the recovery of heterozygous null animals (+/-) following beta-actin-Cre expression. **(B)** Genotyping PCR showing the recovery of wild-type (+/+), heterozygous (+/-), and *Vaultrc5*-null (-/-) animals from *Vaultrc5^+/-^* intercrosses. **(C)** Relative levels of *Vaultrc5* in mouse embryonic fibroblasts isolated from wild-type, heterozygous, and homozygous null animals. **(D)** Top, schematic representation of the *Vaultrc5* locus and its distance to its conserved up- and downstream neighbors. Bottom, expression of *Vaultrc5* flanking genes in brains from wild-type and null animals. **(E)** Observed and Expected numbers for wild-type (+/+), heterozygous (+/-), and Vaultrc5-null (-/-) animals obtained from *Vaultrc5^+/-^* intercrosses. P-value was calculated with a Chi-square test. **(F)** As in E but for animal sex.

### Constitutive loss of *Vautrc5* is compatible with mouse development

To test the broad requirement for vRNA expression during mouse development, we crossed *Vaultrc5*^flx/+^ mice to the general beta-actin-Cre deleter line^24^. This led to widespread recombination of the floxed alleles and to offspring that were heterozygous for a *Vaultrc5* null allele (*Vaultrc5*^-/+^; **Figure 2A**). Subsequent intercrossing of heterozygous *Vaultrc5*^-/+^ mice generated full *Vaultrc5* loss-of-function animals, carrying the null allele in homozygosity (*Vaultrc5*^-/-^) (**Figure 2B**). Quantification of *Vaultrc5* in cells from these animals by RT-qPCR confirmed a genotype-dependent reduction of *Vaultrc5* RNA levels with no detectable transcripts in *Vaultrc5*^-/-^ cells (**Figure 2C**). In contrast, the expression of *Zmat2* and *Pcdha1* was not altered in these samples (**Figure 2D**), confirming that our targeting strategy did not disrupt their regulation.

The recovery of *Vaultrc5*^-/-^ animals at weaning suggests that complete loss of vRNA expression in mice is compatible with animal development. Nevertheless, many genes that have well established essential functions during embryogenesis —including those in the canonical miRNA pathway—lead to lethal phenotypes with incomplete penetrance when disrupted^39–42^. We found no evidence of that being the case for *Vaultrc5*. Specifically, data collected over multiple litters showed that *Vaultrc5*^-/-^ animals were recovered at the expected ratio at weaning (**Figure 2E**). Similarly, we found no deviation from the expected ratios between sexes (**Figure 2F**). Together, these data indicate that mice are able to successfully complete development and survive in the absence of vRNAs.

### *Vaultrc5-*null mice are histologically normal

Aside from being viable, *Vaultrc5*^-/-^ animals were morphologically indistinguishable from littermate controls suggesting that constitutive absence of vRNAs does not lead to gross phenotypic abnormalities in mice. Furthermore, we found no weight differences—a common indicator of suboptimal animal health—between genotypes either in male (median weights of 18.25g, 18.05g, 18.55g for wild-type, heterozygous, and homozygous respectively) or female animals (median weights of 15.75g, 15.9g, 15.8g for wild-type, heterozygous, and homozygous respectively) (**Figure 3A**).

**Figure 3.**
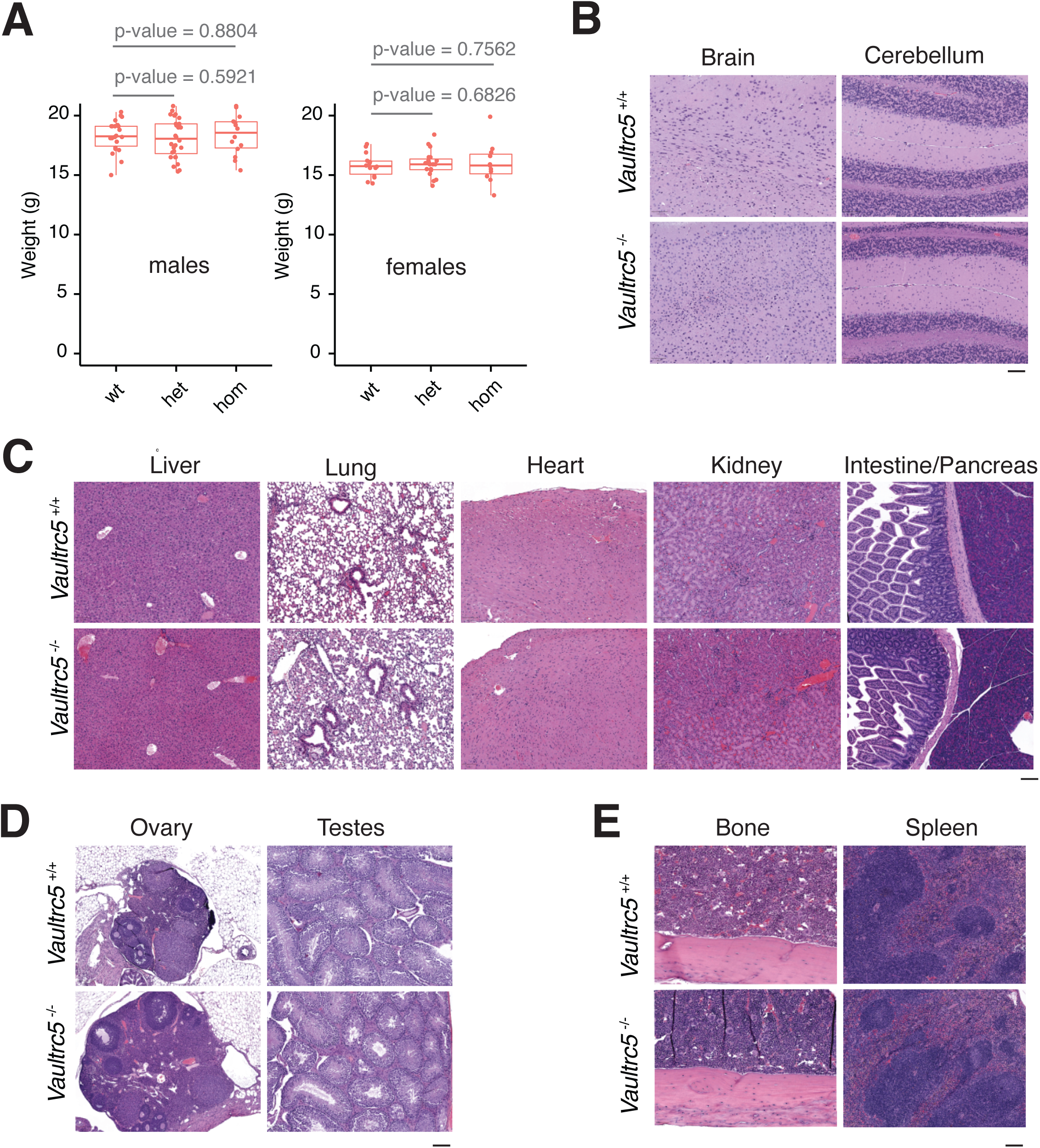
*Vaultrc5*-null mice are morphologically and histologically indistinguishable from wild-type littermates. **(A)** Weight of male (left) and female (right) animals of the indicated genotypes at weaning, showing no difference between *Vaultrc5*-null mice and littermate controls. Each dot represents an animal. p-values were calculated with a two-tailed t-test. **(B-E)** hematoxylin & Eosin-stained histology sections from organs collected from *Vautrc5^+/+^*and *Vautrc5^-/-^* sex matched littermates at 8 weeks of age.

To test if loss of *Vaultrc5* caused more subtle phenotypes, we collected tissues from *Vaultrc5*^-/-^ and *Vaultrc5*^+/+^ littermates of both sexes at 8 weeks of age and subjected them to a histopathological evaluation (**Figure 3B-4E**, **Supplementary Note 1**). We were particularly interested in brain tissues as previous studies had implicated vRNAs in neuronal differentiation^43,44^. In line with a potential role for vRNAs in the brain, their dysregulation has also been implicated in neurodegenerative diseases^45^. Despite these reports, we found that in the absence of *Vaultrc5*, the neural tissues were seemingly uncompromised. Specifically, cortex, medulla, hippocampus, brain stem, corpus callosum and cerebellum all showed no histological differences between *Vaultrc5*^-/-^ and *Vaultrc5*^+/+^ animals (**Figure 3B**). Meninges were present on the cortex, and the parenchyma of the grey and white matter consisted of fine capillaries, glial cells, neurons, and abundant neuropil. The cerebellum of mutant animals was equally unremarkable and characterized by a very cellular granular layer and a less cellular molecular layer, with the prominent Purkinje cells at the interface of the two layers. Finally, the delicate pia mater lined the cerebellum along the molecular layer in animals of both genotypes. In sum, we found no evidence that constitutive loss of vRNA impacts neural tissues in mice, with the histologic findings in both *Vaultrc5*-null and wild-type controls being within normal limits and consistent with normal brain development and tissue differentiation.

**Figure 4.**
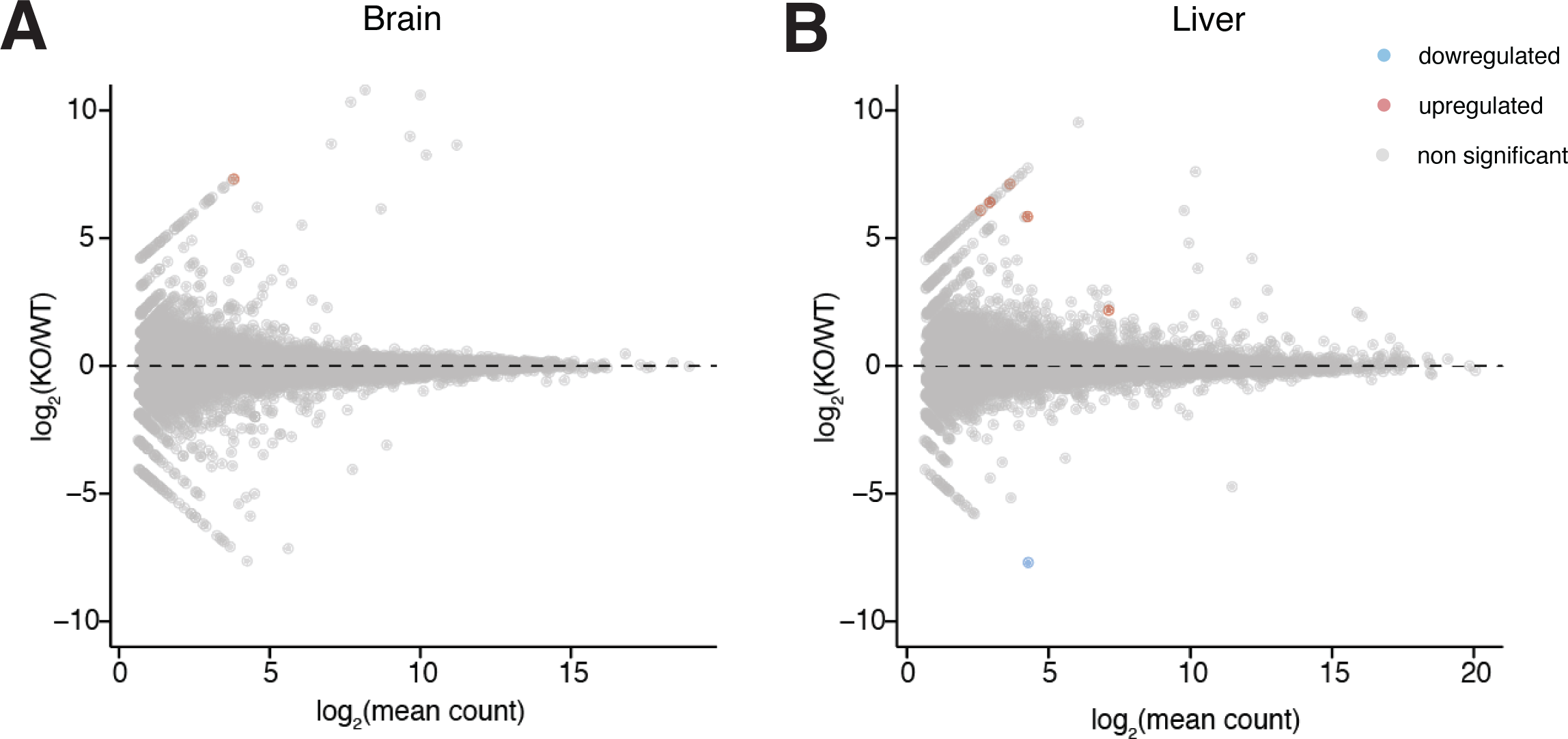
RNA sequencing profiles of *Vautrc^+/+^* and *Vautrc5^-/-^* animals. MA plots showing lack of differentially expressed genes between brains **(A)** and livers **(B)** of wild-type and Vaultrc5-null animals. Differentially expressed genes were defined as those with log_2_ fold- change > [2] and adjusted p value < 0.1.

In addition to neural tissues, *Vaultrc5* may also impact liver biology since vRNAs were initially identified as components of the Vault particle from rat liver extracts^1^, an organ in which the murine *Vaultrc5* is well expressed^32^. Moreover, this organ is characterized by high levels of autophagy^46^, a process that seems to be regulated by vRNAs through their interaction with p62 via the central region (**Figure 1B**) ^19^. Although the predicted structure of this region has limited similarity between *vtRNA1-1* and *Vaultrc5* (**Figure 1A**), binding of p62 to vault RNAs seems to be conserved between the two species ^19^. Yet, despite the importance of autophagy to hepatic functions^46^, histopathological analysis showed no evidence that the development or structure of the liver was compromised in the absence of *Vaultrc5*. Specifically, the liver in animals of both genotypes was encapsulated and composed of hepatocytes with abundant granular pink cytoplasm. In both cases the organ was organized as cords of cells with a zonal arrangement around both portal triads and central veins. Macrophages and scattered mononuclear cells were also identified throughout the parenchyma (**Figure 3C**). Similarly, in our analysis of other major organs including the lung, heart, kidney, spleen, intestine, pancreas (**Figure 3C**) as well as ovaries, and testis (**Figure 3D**) we found no evidence of histological differences between *Vaultrc5*^-/-^ and *Vaultrc5*^+/+^ animals. A detailed description of the histological findings in these organs can be found in **Supplementary Note 1**.

Finally, we analyzed the histology of bone and spleen sections in our mice as these are major hematopoietic and lymphoid organs and vRNAs have been previously implicated in immune cell functions^47–49^. We found no histological differences between the spleens of *Vaultrc5* wild-type and *Vaultrc5* loss-of-function animals (**Figure 3E**). Similarly, bones of wild-type and mutant animals were histologically identical (**Figure 3E**). In both cases bone marrows showed high cellularity and tri-lineage hematopoiesis. Within this cell population, megakaryocytes were the most abundant cells with erythroid and granulocytic precursors easily observed at higher magnifications. In all cases these cells showed no morphological abnormalities in the absence of *Vaultrc5*.

As part of our pathology analysis, we also performed a complete blood count. We found that measurements for *Vaultrc5*^-/-^ animals as well as for *Vaultrc5*^+/+^ littermate controls were within the normal physiological ranges expected for the C57Bl/6 strain (Mouse Phenome Database, www.jax.org/phenome). They were also for the most part identical between wild-type and mutant mice with two notable exceptions (**Supplementary Figure 3**). First, although we found no statistical difference in white and red blood cell counts (**Supplementary Figures 3A and 3B**), red blood cell size (**Supplementary Figure 3E, 3F**), or hemoglobin levels (**Supplementary Figure 3G-I**) between *Vaultrc5*^-/-^ and *Vaultrc5*^+/+^ animals, hematocrit values—the percentage of red blood cells in the blood—were slightly reduced in mutant mice (**Supplementary Figure 3D**). This suggests that *Vaultrc5* may have essential albeit subtle roles in erythropoiesis. Second, the platelet counts were reduced in the absence of *Vaultrc5* to approximately 78% of the values in wild-type animals (**Supplementary Figure 3C**). This may point to a potential role for *Vaultrc5* in platelet development from megakaryocyte precursors in the bone marrow or in the regulation of platelet survival. Nevertheless, platelet values remained well within the normal physiological ranges expected for the C57Bl/6 strain (*Vaultrc5*^+/+^=977-997 x 10^3^ platelets/μl; *vautrc5*^-/-^=745-792 x 10^3^ platelets/μl; C57Bl/6=562-2159 x 10^3^ platelets/μl).

### RNA sequencing profiles of wild-type and *Vaultrc5^-/-^* animals

We reasoned that even in the absence of major histological changes, loss of *Vaultrc5* could result in molecular phenotypes that would be reflected in transcriptome dysregulation. To determine if that was the case we performed RNA sequencing to the brain and liver of *vautrc5*^-/-^ animals, alongside those of age- and sex-matched wild-type littermate controls (**Figure 4**). As before, we chose to focus on these organs because previous studies pointed to a role for vRNA in their development or homeostasis. For example, *in vitro* studies both human *vtRNA1-1*^43^ and mouse *Vaultrc5*^44^ promote synapse formation by modulating MAPK signaling and deregulation of the human *vtRNA2-1* was proposed to induce neural dysfunction in humans^45^. Despite this, we found no evidence of gene dysregulation caused by absence of vRNA in mouse brains (**Figure 4A**). Similarly, human *vtRNA1-1* was shown to promote cell proliferation and tumorigenesis in human liver cell lines^37^, which together with the abundant expression of vRNAs in the liver^32^ suggests a potential role for vault RNAs in this organ. Yet, we again found no gene significantly dysregulated in the *Vaultrc5*^-/-^ samples (**Figure 4B**). Together with the histological analysis to these organs, these data suggest that in the mouse, vRNA has no significant impact to the development or homeostasis of the brain or liver.

## DISCUSSION

We describe here the generation of *Vaultrc5* conditional knockout mice and the initial characterization of animals with constitutive deletion of this locus. Given that mice—like most mammals—have a single *vtRNA1* gene and have in addition lost *vtRNA2* ^23^, *Vaultrc5*^-/-^ animals are null for vRNAs. This is to our knowledge the first characterization of the physiological requirements for this evolutionarily conserved but poorly understood class of small RNAs in vertebrates.

Despite the myriad of roles assigned to vRNA genes in humans, ranging from regulation of cell proliferation, survival, and differentiation, we found that animals null for *Vaultrc5* are viable and histologically normal. This suggests that *vRNAs* may not be essential in mice under unchallenged conditions. This is similar to what has been described for the other components of the Vault particle. Indeed, mice null for the Major Vault Protein (*Mvp*^-/-^), the telomerase-associated protein 1 (*Tep1*^-/-^), or the vault poly-(adenosine diphosphate-ribose) polymerase (*Vparp*^-/-^) are all seemingly phenotypically normal^50–52^ even when multiple genes are deleted^52^. Yet, when challenged with specific pathogens *Mvp*^-/-^ animals show increased resistance to infection^53^. Similarly, *Varp*^-/-^ animals show increased susceptibility to carcinogen induced tumorigenesis^54^. Thus, as is the case for many other genes, it may be that *vRNA* expression only becomes indispensable under particular conditions such as aging, oncogenic stress, or viral infections all of which have been previously linked to *vRNA* functions. Hematopoiesis may be particularly sensitive to loss of *vRNAs* since we have observed a small but significant reduction in the platelet counts in our animals compared to littermate controls. Follow-up studies breeding the conditional *Vaultrc5*^fx/flx^ animals with lineage restricted deleters such as *Vav-Cre* (leading to recombination of the locus in all cells of the hematopoietic system)^55^ or *Gp1ba-Cre* (leading to recombination in megakaryocytes)^56^ combined with both immune challenges and a more detailed characterization of the immune system will be essential to fully characterize these phenotypes, define the extent to which *Vautrc5* is required for hematopoiesis, and understand the predominant mechanism through which this evolutionarily conserved small non-coding RNA regulates mammalian physiology. To facilitate those studies, we are making our mice freely available to the community through *The Jackson Laboratory* mouse strain repository.

## Supporting information

SupFigure1

SupFigure2

SupFigure3

SupNote1

SupTable1

## Acknowledgments

We thank all members of the Vidigal and Batista labs for discussions and comments on this work. We thank Raj Chari and Parirokh Awasthi for the generation of the *Vaultrc5* allele. This work was supported by the Intramural Research Program of the National Institutes of Health through the Center for Cancer Research, National Cancer Institute, project 1ZIABC011810 (J.A.V.)

## Figure Legends

**Supplementary Figure 1. Conservation and targeting of the mouse *Vaultrc5* locus. (A)** Genome browser view of the syntenic region containing the *Vaultrc5* and its conserved upstream neighbor *Zmat2*. PhyloP basewise conservation scores for Placental mammals and for Vertebrates, as well as UCSC Refseq gene annotations are shown. **(B)** As in (A) but zoomed in on *Vaultrc5* locus. Vertical doted lines highlight the position of the inserted *loxP* sites in the conditional allele. Position of known regulatory elements are shown below. **(C)** sequence alignment of the conserved regions between *Vaultrc5* and the three human vtRNA1 genes which overlap the A and B2 boxes (in green) and the termination sequence (in red). Dots highlight the nucleotides that differ between the murine gene and at least one of the human vtRNA1 genes. Grey boxes highlight the nucleotides that differ between *Vaultrc5* and *vtRNA1-1*. **(D)** Sanger sequencing of the integrated donor template showing the position of the loxP sites, the XmnI restriction sites, and the PAMs of the gRNAs used.

**Supplementary Figure 2. Analysis of putative miRNAs derived from the *Vaultrc5* locus. (A)** Comparison of the predicted human and mouse vaultRNA-derived miRNAs. The seed-sequence is highlighted in bold. Nucleotides that differ between the two species are highlighted in pink **(B)** Genome browser view of sequencing reads over *Vaultrc5* and *pre-mmu-miR-19a* loci in multiple adult mouse tissues in triplicate sequencing experiments **(C)** Abundance of reads (as RPM) for predicted vaultRNA-derived fragments. Error bars represent standard deviation between triplicate experiments. **(D)** As in (C) but for *miR-19a-3p* and *miR-19a-5p*.

**Supplementary Figure 3. Complete blood counts for *Vautrc5^-/-^* and *Vautrc5^-/-^* animals.** Cell count for White blood cells **(A)**, Red blood cells **(B)**, Platelets, **(C)**. Assessed hemoglobin levels by Hematocrit **(D)**, Mean Red Blood Cell volume **(E)**, Red Blood Cell distribution width **(F)**, Hemoglobin **(G)**, Mean cell hemoglobin **(H)**, Mean cell hemoglobin concentration **(I)**. Error bars represent standard deviation between biological replicates. p-values were calculated with a t-test and are highlighted in green when below the significance threshold of 0.05.

