## Supplementary figures and images for "The small non-coding RNA *Vaultrc5* is dispensable to mouse development"

### SupFigure1

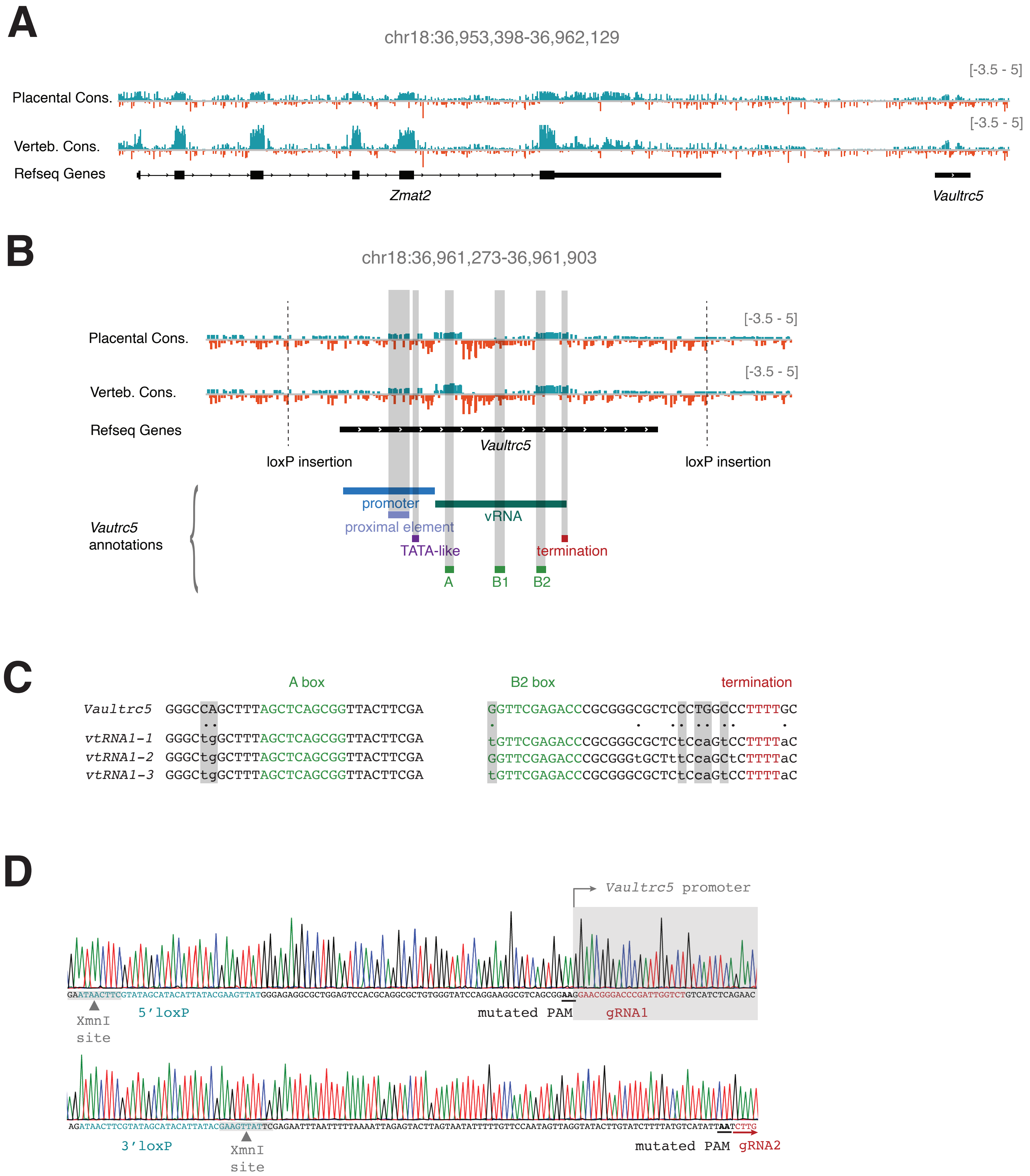


**Prajapat Supplementary Figure 1**

### SupFigure2

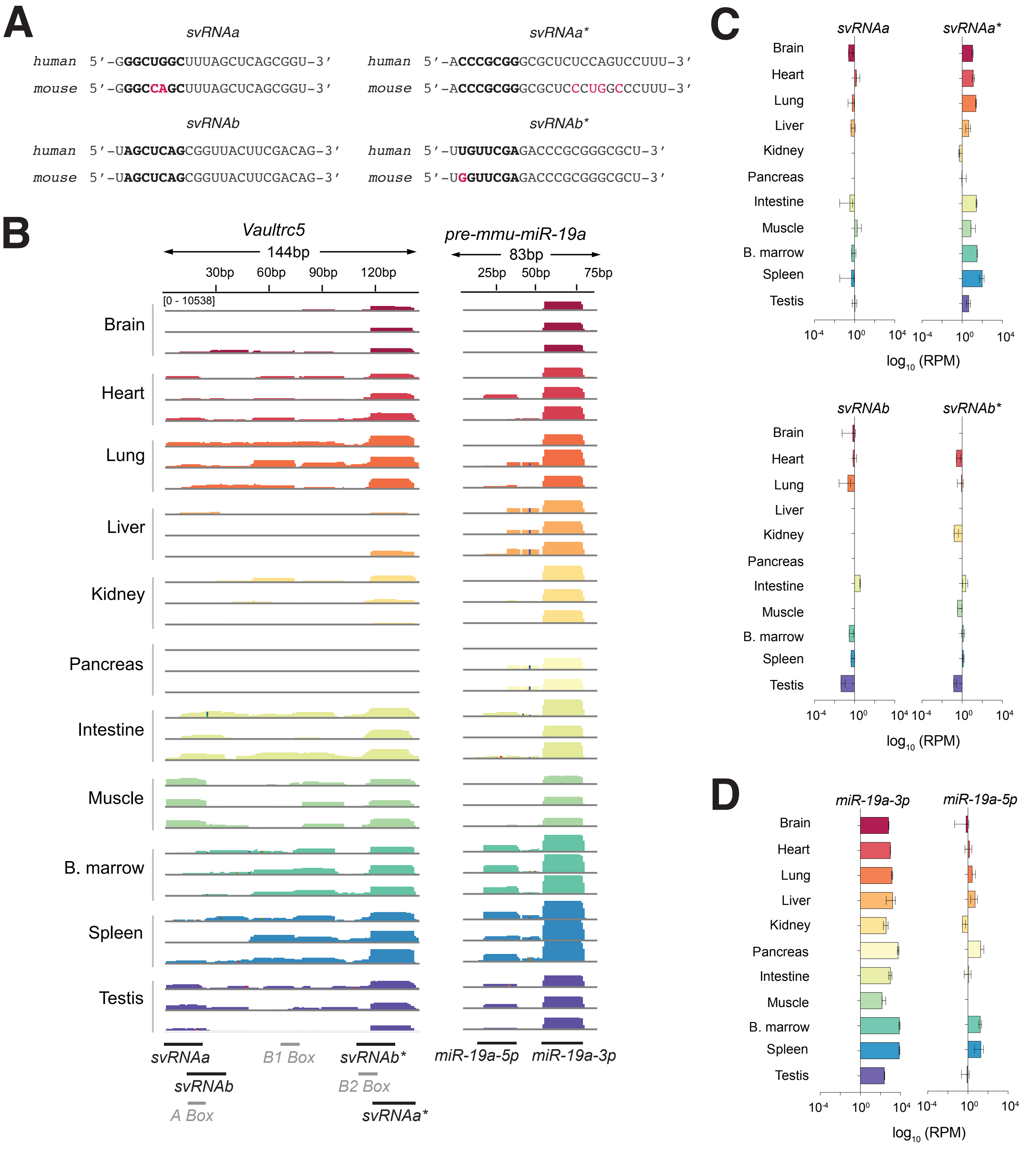


**Prajapat Supplementary Figure 2**

### SupFigure3

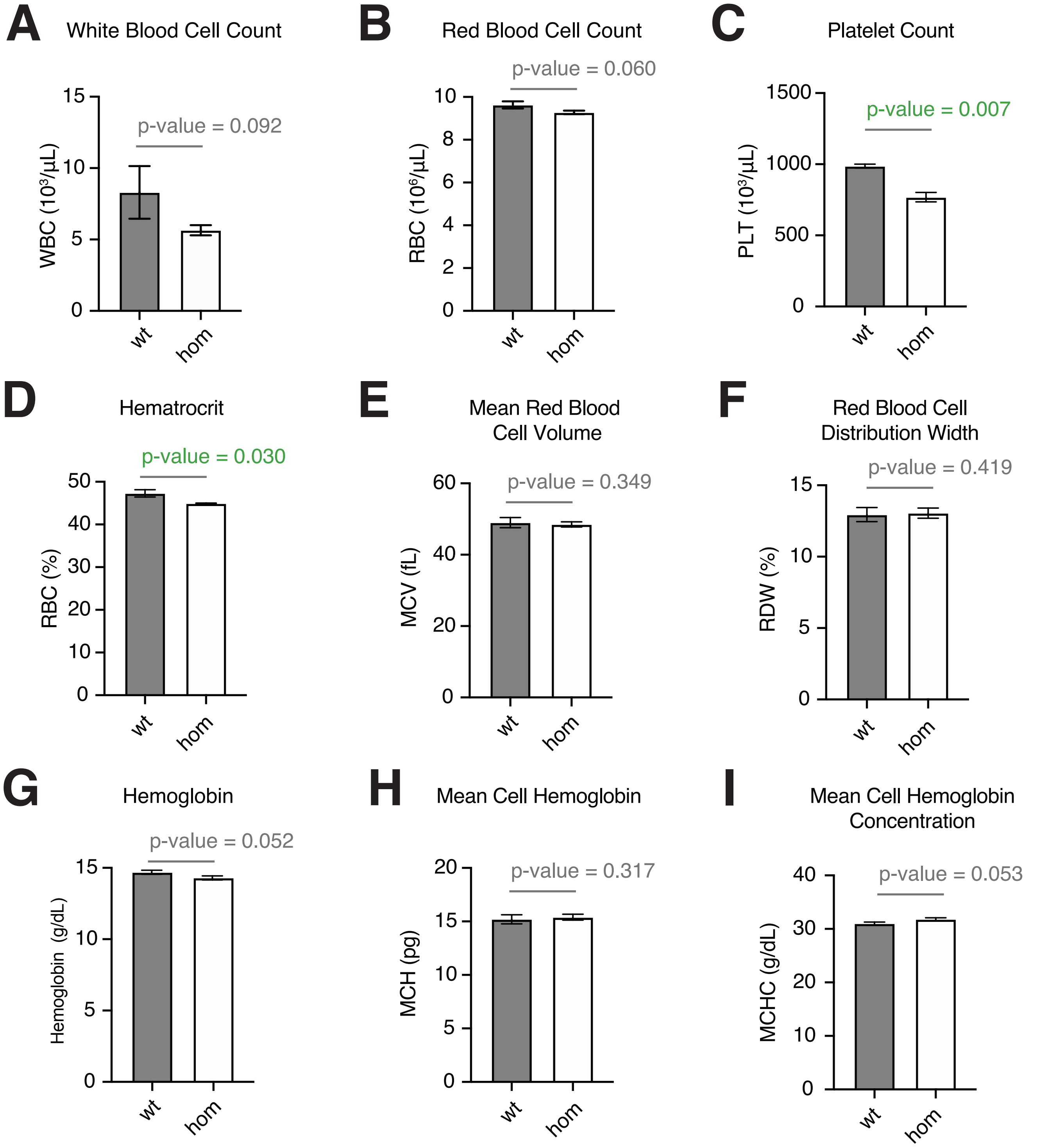


**Prajapat Supplementary Figure 3**
