## Supplementary material for "The small non-coding RNA *Vaultrc5* is dispensable to mouse development": SupNote1

**Supplementary Note 1**

Additional histological findings in *Vaultrc5^-/-^* versus *Vaultrc5^+/+^* animals:

Lung: The lung parenchyma of *Vaultrc5^-/-^* animals was unremarkable. Both bronchi and bronchioles were lined by columnar epithelium. Arteries and veins were also easily identified adjacent to large and medium-sized airways within a loose connective tissue stroma. Alveoli were thin walled and attenuated by pneumocytes with red blood cells interspersed within very small capillaries. The alveolar spaces were empty, as expected.

Heart: The heart of loss-of-function mutants like the heart of wild-type animals was lined by endocardium. In both cases the pericardium could be identified on the outer surface of the organ. The chambers are unremarkable and contained blood. The myocardium was composed of myocytes with a single, central nuclei. Small and very small blood vessels/capillaries could be identified by the presence of red blood cells between the myocardial fibers. At the superior aspect, adipose tissue containing cardiac vessels, pulmonary vessels and occasionally bronchial tissue was present, all of which were unremarkable where observed.

Kidney: For both male and female animals of both genotypes the kidneys were encapsulated and composed of an outer cortex layer and an inner medulla, each with a distinct cortico-medullary junction. The glomeruli were interspersed in the cortex, and the collecting duct system could be identified in the medulla. The tubules were normally formed, and blood vessels as well as the interstitium were in all cases unremarkable. The hilar and perirenal adipose tissue were equally unremarkable, and lymph nodes, as well as vessels, could be identified within the hilar tissue.

Intestine and Pancreas: *Vaultrc5^-/-^* animals showed normal intestinal architecture with villi and crypts composed of intestinal epithelium interspersed with goblet cells. The interstitium contained a normal number of inflammatory cells, and the muscularis mucosae, submucosa and muscularis propria are unremarkable. No histological differences were detected when compared with wild-type controls. These same gastrointestinal sections also contained unremarkable pancreas tissue. Specifically, the parenchyma was composed of acini tissue (exocrine), small pancreatic ductules as well as larger ducts, blood vessels and interstitium. Islets of Langerhans (endocrine) were also observed in both mutant animals and controls.

Ovaries and testis: We found no histological differences between the ovaries of wild-type and *Vaultrc5*-null females. In both cases, they were composed of an outer cortex and central medulla. Follicles of various stages were surrounded by cortical stroma and characterized by a granular cell layer of cuboidal epithelium and a central clearing. No definitive oocytes were observed. Corpora lutea were also prominently seen. Adjacent fallopian tube tissue had a normal cuboidal epithelium, and adjacent adipose and fibrovascular tissue was also unremarkable. The testes were also unremarkable, with seminiferous tubules and epididymis observed in both genotypes. The seminiferous tubules contained numerous germ cells, inconspicuous Sertoli cells, and were in both cases attached to a thin basement membrane. In the lumen, normal spermatogenesis was observed. The epididymis had smooth muscle, pseudostratified, ciliated columnar epithelium, and a basal cell layer between the two. Sperm are present in the lumen.
